# Combining formal education and citizen science: A case study on students’ perceptions of learning and interest in an urban rat project

**DOI:** 10.1101/2020.01.27.921395

**Authors:** Tuomas Aivelo, Suvi Huovelin

**Affiliations:** Organismal and Evolutionary Biology research program, University of Helsinki

**Keywords:** citizen science, near environment, urban ecology, animal studies, animal attitudes

## Abstract

Citizen science is a valuable tool in environmental and formal education in creating scientific knowledge for the researchers and facilitating learning and fostering a positive relationship toward the environment and study species. We present a case study on the Helsinki Urban Rat Project in which students surveyed rat occurrence in their own near environments. According to our results, experientiality, involvement, meaningfulness, freedom to choose, ease of participation, and the rats themselves contributed to students’ increased interest in participation. Furthermore, students described diverse factual, conceptual, procedural, and metacognitive knowledge that they acquired during their participation. In general, students described negative attitudes toward rats, but they described fewer negative views on rats after participation. We reflect on the success of the citizen science project and implications of planning a future citizen science project and incorporating citizen science in formal education.

## Introduction

Public participation in science has a long history and recently there has been a renewed interest through a concept called *citizen science* (Miller-Rushing, Primack, and Bonney 2012; Silvertown 2009). Citizen science is defined as systematic research done by non-professional researchers with co-operation from professional researchers (Dickinson et al. 2012). Citizen science can involve lifelong participation and sophisticated research skills, or it can be short-term with no specific skills required to participate (Shirk et al. 2012). Public participation in science has been suggested to lead to a better understanding of science and scientific processes and foster positive attitudes towards science (Ruiz-Mallén et al. 2016; Phillips et al. 2018; Bonney et al. 2016). Furthermore, the research literature suggests that doing research in one’s own *near environment* (i.e., environment in which a person lives or works) nurtures positive attitudes toward the environment, and specifically the studied species (McKinley et al. 2017; Ballard, Dixon, and Harris 2017). Citizen science is adopted in formal science education as authentic research is seen as motivational to students (Zoellick, Nelson, and Schauffler 2012; Trumbull et al. 2000; Hofstein, Eilks, and Bybee 2011). Nevertheless, there is a lack of evidence about how students learn and how their attitudes are shaped in authentic citizen science projects.

We present a case study on the citizen science dimension of the Helsinki Urban Rat Project (HURP), which investigates how participating upper- and lower-secondary students perceive learning about urban rats. We seek to answer the following research questions:

1. What aspects do the students perceive as interesting or uninteresting in the study?
2. What aspects of learning during the project do the students describe?
3. How are rats perceived as a focus of a citizen science project?

### Citizen science and environmental education

#### Participating in doing science

In environmental research, citizen science has long been used in various initiatives, such as bird ringing and butterfly and weather data collecting (Miller-Rushing, Primack, and Bonney 2012), and it has encountered a renewed interest in recent decades because there is widening public participation in science. The main tenet of citizen science is to allow the wider public to participate in the scientific process (Bonney et al. 2016). In a democratic society, this is seen as a part of the public’s right not only to knowledge, but also to knowledge-creating processes.

Citizen science projects can be classified in three distinct categories: contributory, collaborative and co-creative (Bonney, Ballard, et al. 2009). Contributory projects are those that benefit from data collected from citizen scientists, and collaborative projects include citizen scientists in the research planning and data analysis. Co-creative projects involve citizen scientists from the very beginning in shaping the research plan and at every step of the research. Most of the citizen science projects belong in the contributory category (Bonney, Ballard, et al. 2009); thus, while there are an increasing number of citizen science initiatives and large swaths of research data have been collected by citizen scientists, the relationship between professional researchers and citizen scientists sometimes seems one-way (Shirk et al. 2012; Rotman et al. 2012). As citizen science projects are planned by professional researchers who prioritize their own needs, there has been unsurprisingly less emphasis on what citizens participating in research can actually learn (Phillips et al. 2018).

#### Relationship between humans and their near environments

Citizen science projects can profit from a voluntary workforce, as well as from knowledge residing in communities, such as knowledge of the near environment (McKinley et al. 2017). Citizen science can make use of experiential knowledge (Smith 2006; Harkness 2007), embodied knowledge (Lawrence and Shapin 1998), or situated knowledge (Haraway 1988) as something that the professional researchers are not capable of producing themselves. Citizen science can be a means to explore a near environment, survey the current status of the environment, and even track changes over a period of years (Pollock and Whitelaw 2005). While citizen science has a number of justifications and objectives, ranging from enhancing public appreciation of science to teaching specific research skills, it is also increasingly used as a tool for environmental education because it has been argued that citizen science can facilitate the emergence of positive relationships with nature (Ballard, Dixon, and Harris 2017).

Sensitization by seeing the effects of human influence in the environment can further make it easier to reflect on environmental change (Krasny and Bonney 2005). As a relationship with the near environment develops throughout human life, positive experiences in nature can foster a willingness to protect the environment and biodiversity (Hosaka, Numata, and Sugimoto 2018). Citizen science can also affirm pre-existing attitudes; for example, worrying about the near environment can prompt active participation in citizen science (Pollock and Whitelaw 2005).

#### Citizen science in schools

Citizen science has certain peculiarities that make some projects challenging. Firstly, participation in a school context is not voluntary, but rather students are compelled to do science as part of their regular curriculum. Thus, participation is usually driven by curriculum and teachers’ decision on how to implement that curriculum. This can affect the students’ interest and learning outcomes during school-mandated citizen science (Shah and Martinez 2016; Rotman et al. 2012). Secondly, as schools have a detailed curricula, citizen science in a school context becomes a part of formal education. This adds additional pressure to citizen science to conform to the institutional practices of schools. This means that care is needed to ensure that citizen science participants’ and researchers’ needs align with school curricula (Zoellick, Nelson, and Schauffler 2012). Thirdly, participating in citizen science in school can drive interest in participating in citizen science voluntarily outside of the school setting (Silva et al. 2016).

In turn, schools provide many opportunities for citizen science; teachers can support students during the process, and schools can sustain long-term commitments to citizen science projects (Dickerson-Lange et al. 2016; Rock and Lauten 1996). Furthermore, citizen science can be successfully incorporated in the curriculum because there is increasing emphasis on inquiry-based learning and out-of-school teaching (Shah and Martinez 2016).

## Theoretical background

To understand how well citizen science projects can facilitate learning in a formal school education context, we studied students by interviewing them in groups about how they perceived participation in a citizen science initiative aimed to study urban rats. To conceptualize students’ descriptions of their learning, interest, and attitudes toward rats, we use educational theories, such as situational and personal interest, Bloom’s taxonomy on learning objectives, and Kellert’s classification of animal attitudes.

### Interest in doing citizen science

*Interest* is one of the main factors driving learning and positive attitudes toward science (Palmer 2004). Interest is a psychological state or selective positive feeling toward a certain subject (Ainley, Hidi, and Berndorff 2002). Interest has been conceptualized as compromising to distinct components: situational interest and personal interest (Hidi and Renninger 2006). Situational interest is a type of interest that is short-term and externally driven, whereas personal interest is long-term and borne out of personal significance. Nevertheless, sustained situational interest can lead to the emergence of personal interest (Hidi and Renninger 2006). Many external factors have been shown to lead to peaking situational interest: novelty (Palmer 2009), discrepant events (Lin, Hong, and Chen 2013), and meaningfulness or involvement (M. Mitchell 1993). Novelty means that observed phenomena are novel for the observer, and discrepant events refer to phenomena that are unusual from the observer’s perspective. Meaningfulness is borne from a connection between participants’ everyday lives and observed phenomena, whereas involvement is created by freedom to act during the research process.

Interest during citizen science participation has not been studied extensively. It is known that citizen science projects can ignite interest in science and sustain learning (Hiller and Kitsantas 2014; Silva et al. 2016); however, it is not clear what aspects of citizen science are important. For example, out-of-school experiences can promote interest in science (Uitto et al. 2006). In our study, we qualitatively surveyed descriptions of interest from students through interviews and content analysis.

### Learning outcomes in citizen science

Because citizen science as an institutional practice is a novel phenomenon, the learning objectives of citizen science projects are usually not well formulated (Phillips et al. 2018). Most often, observed learning outcomes are on the level of factual knowledge on the study species, environment, or phenomena (Jordan et al. 2011; Silva et al. 2016; N. Mitchell et al. 2017). Nevertheless, participants also acquire procedural knowledge on doing research in general and with specific research methods (Trumbull et al. 2000), though there are also contrary results (Brossard, Lewenstein, and Bonney 2005). Many researchers claim there is limited evidence of an increase in a deeper understanding of the scientific processes (Bela et al. 2016; Bonney et al. 2016; Jordan, Ballard, and Phillips 2012). It is significant that citizen science participation can also lead to affective learning, such as changes in attitudes toward environmental problems (McKinley et al. 2017).

In recent years there has been an effort to define the educational outcomes of citizen science projects (Phillips et al. 2018). The basis for this work has been informal science education frameworks such as one outlined by Allen et al. (2008). As we studied participation in citizen science in a formal education context, the widely used revised Bloom’s taxonomy (Krathwohl 2002; Bloom et al. 1956) was a suitable framework to assess learning outcomes. The revised Bloom’s taxonomy divides learning objectives in a two-dimensional matrix with a cognitive process dimension classifying increasingly complex cognitive tasks and a knowledge dimension, which is a range from concrete to abstract processes. While Bloom’s taxonomy has been criticized for its behaviorist underpinnings, and it does not form an optimal model of learning in citizen science projects, it did provide a suitable framework to classify and analyze learning outcomes that students have described in our study.

### Citizen science and attitudes about animals

Citizen science fosters human-animal relationships by making the presence and absence of animals more tangible (McKinley et al. 2017). In a school context, contact with animals and nature have been shown to foster a positive relationship toward nature (Fox-Parrish and Jurin 2008; White, Eberstein, and Scott 2018; Palmberg and Kuru 2000). In general, children are more prone to explain a need to protect species that they describe as pleasing in comparison with less agreeable species (Almeida, Vasconcelos, and Strecht-Ribeiro 2014). Concerning disliked or “disgusting” species, it is not yet established how likely studying them will decrease negative attitudes. For example, studying living snails leads to higher knowledge scores and less disgust sensitivity (Prokop and Fančovičová 2017), but in comparison, when students studied strongly disliked prairie dogs, there was no positive change in students’ attitudes (Fox-Parrish and Jurin 2008).

Based on a large sample of questionnaire data, Kellert (1985) suggested 12 factors that are important for public preferences for different animal species: size, aesthetics, intelligence, danger to humans, likelihood of inflicting property damage, predatory tendencies, phylogenetic relatedness to humans, cultural and historical relationship, relationship to human society, texture, mode of locomotion, and economic value of the species. Thus, humans tend to like animals that are economically or emotionally important, phylogenetically or habitually like humans, large-sized, intelligent, or esthetically pleasing. Humans dislike animals that are dangerous, pathogen vectors (Curtis, Aunger, and Rabie 2004; Löe and Röskaft 2009), pests (Breitenmoser Urs 1998), or different looking than humans (Almeida, Vasconcelos, and Strecht-Ribeiro 2014).

## Our aims and research questions

We conducted group interviews with students to understand how the students described their interest in doing citizen science and their learning. As the study animal, the brown rat (*Rattus norvegicus*) is generally perceived as a pest, and is one of the most disliked animals in urban environments. We were interested in understanding how the students perceived the rat as a study subject. Our aim was to provide more information on how well citizen science projects and formal learning in a school context can be combined.

Specifically, we had the following research questions:

1. What aspects of the citizen science project are described as interesting or uninteresting?
2. What facts, concepts, or skills do the students perceive they have learned during the project?
3. How do students reflect on their perceptions of rats and how has studying rats affected that relationship?

## Materials and methods

We performed a qualitative case study on student perceptions of citizen science participation and studied wild rats in the Helsinki Urban Rat Project by conducting group interviews and analyzing those interviews through an iterative, theory-guided content analysis.

### Helsinki Urban Rat Project and track plates

The context in which this study took place was the Helsinki Urban Rat Project (HURP; https://www.helsinki.fi/en/projects/urban-rats), an ongoing multidisciplinary research project. HURP aims to understand how rat populations vary spatiotemporally in the urban landscape, how parasites and pathogens are spread in rat populations and in human-rat interfaces, and how humans perceive sharing the same habitat with invasive rats. The study organism in this citizen science project is the brown rat (*Rattus norvegicus*), which usually elicits strong negative reactions from humans. Attitudes toward rats are very negative globally (Bjerke and Østdahl 2004; Kellert 1985; Collins 1976; George et al. 2016). Arrindell (2000) classified the rat as a “fear relevant animal” because the rat causes more fear than actual danger. Based on the preliminary results of HURP, rats are common urban animals throughout the study area (unpubl.). The citizen science part of HURP consists of collecting presence-absence data on urban brown rats. The rat occurrence data are collected with track plates (Hacker et al. 2016), which are 20×20-cm white plastic plates that are painted with a lampblack-ethanol mixture. When this paint dries, it leaves a black surface that flakes away when an animal steps on the plate, thus showing paw prints for any animals walking on the plate. Rat tracks can be easily distinguished from other urban animals because of their characteristic shape and size.

We recruited lower- and upper-secondary school biology teachers and offered them a task as part of the science project, which was aimed to fulfill curricular demands in both lower- and upper-secondary school. The participation began when the first author visited the classroom, gave a lecture on urban ecology and brown rats and explained how to participate in the research project. Teachers were free to organize the participation in the research however it best fit their teaching. The minimum requirements were that four plates were left in one site, kept there for four days, and photographed daily. Furthermore, when sending the research data, students described the overall environment where the plates were left and counted the rat tracks on the plates. The students sent data to us through the Epicollect5 mobile application (Aanensen et al. 2009). In general, the students were free to decide where they set the plates, and they were encouraged to study environments with which they were familiar, such as near their homes, schools, or places where they otherwise spend time. We asked students to place the track plates in places where rats are likely to move, so students were compelled to reflect on their study area from the point of view of rats and how they use space.

### Student interviews

We recruited students for interviews from the classes that had already participated in the HURP citizen science project during the past month. We interviewed 2–4 students at a time semi-structurally in a group setting from October 2018 to January 2019. The selection of interviewees was based on volunteers. There was a total of 29 interviewees, of which 14 were from lower-secondary school (4 girls and 10 boys whose ages ranged between 15 and 16 years) and 15 from upper-secondary school (10 girls and 5 boys with an unknown age range but likely between 16 and 19 years). The length of interviews was from 12 minutes to 31 minutes, with a mean length of 25 minutes.

We chose a group interview as the data collection format because it allowed for interviewing more students and it facilitated the reflection of the learning process among young people who might not have been as reflective when interviewed individually. The interviews contained two main themes: 1) interest and perceived learning and 2) attitudes toward rats. They were conducted in the school during regular classroom times.

### Ethical considerations

The research permit was granted by the City of Helsinki on April 5, 2018, and the permit from an individual private school on October 1, 2018. The requirements of the research permit included permission from both the principal and the biology teacher to ask students to participate. All participants were over 15-years old so, which placed them under the guidelines from the City of Helsinki and Finnish National Board of Research Integrity TENK and no permissions were required from their guardians. The parents were informed prior to the interview by electronic letter. All participants were informed about the aim of the study and how the materials would be collected, stored, and handled anonymously. It was made clear that participation was voluntary, participation could be ended at any time, no data would be given to their teachers, and participation would not affect their grades.

### Qualitative analysis

We performed an iterative, theory-guided content analysis. We started transcribing interviews after the very first interviews and we continued doing it concurrently as we performed further interviews. In an iterative fashion, after each interview we refined the interview frame by modifying questions and choosing expressions that were understood by most students. We also began to analyze the data as soon as the first interviews were transcribed.

The analysis units in the transcribed texts varied from a full sentence to individual words. During the analysis, the units were simplified, classified and grouped. As we had no prior experience on how students react or what they perceive to learn during the project, the interviews were quite exploratory. Thus, we had to use three different theoretical frameworks to analyze the interviews to provide a rich interpretation of student experiences during their participation.

Firstly, as the theoretical framework for student interest, we used the division of situational and personal interests (Ainley, Hidi, and Berndorff 2002). Secondly, for the analysis of learning outcomes, we classified student interviews with the revised Bloom’s taxonomy (Krathwohl 2002). Thirdly, for student descriptions of attitudes toward animals, the theoretical background used was Kellert’s (1985) classification of feelings directed toward animals. Through themes and types, original student comments were compared to the theoretical background and disconfirming data were highlighted and new categories were created when needed.

### Trustworthiness of the study

The credibility of the study was enhanced by using interviews as the data collection method. Interviews allowed for asking further questions to ensure that the student meanings were understood correctly. Furthermore, interviewing provided direct evidence of the students’ perceptions of the project. The selection of interviewees was based on which students volunteered, which could bias what experiences of participants were heard by researchers. Nevertheless, the school groups were small; the groups consisted of students who worked together with other students in doing the track plates surveys, and they had broadly similar experiences. Transferability is a challenging aspect of this study because it is a case study based on a unique project. We kept a detailed audit trail of the research context, data collection, and analysis to make reporting as accurate and detailed as possible. Confirmability was established by the audit trail and taking into account researcher positionality and bias. Both researchers confess to having positive attitudes towards rats, and this was considered during the analysis of attitudes toward rats. The first author is the principal investigator in HURP and is leading the citizen science project. Thus, he encouraged critical handling of the project during the analysis phase. In relation to the dependability, the analysis was conducted by the second author. After creation of the classifications in dialogue with the first author, the first author repeated the analysis to measure inter-rater reliability. The inter-rater reliability was measured with Cohen’s Kappa (Cohen 1960). In all classifications it was 0.81, which suggests excellent agreement.

## Findings

### Interest and disinterest in rat research

In every interview at least one interviewee said that participation in the project was interesting. This interest was classified in 10 subclasses that were in turn grouped to 6 main classes, which were divided in two types of interest (Table 1).

**Table 1:**
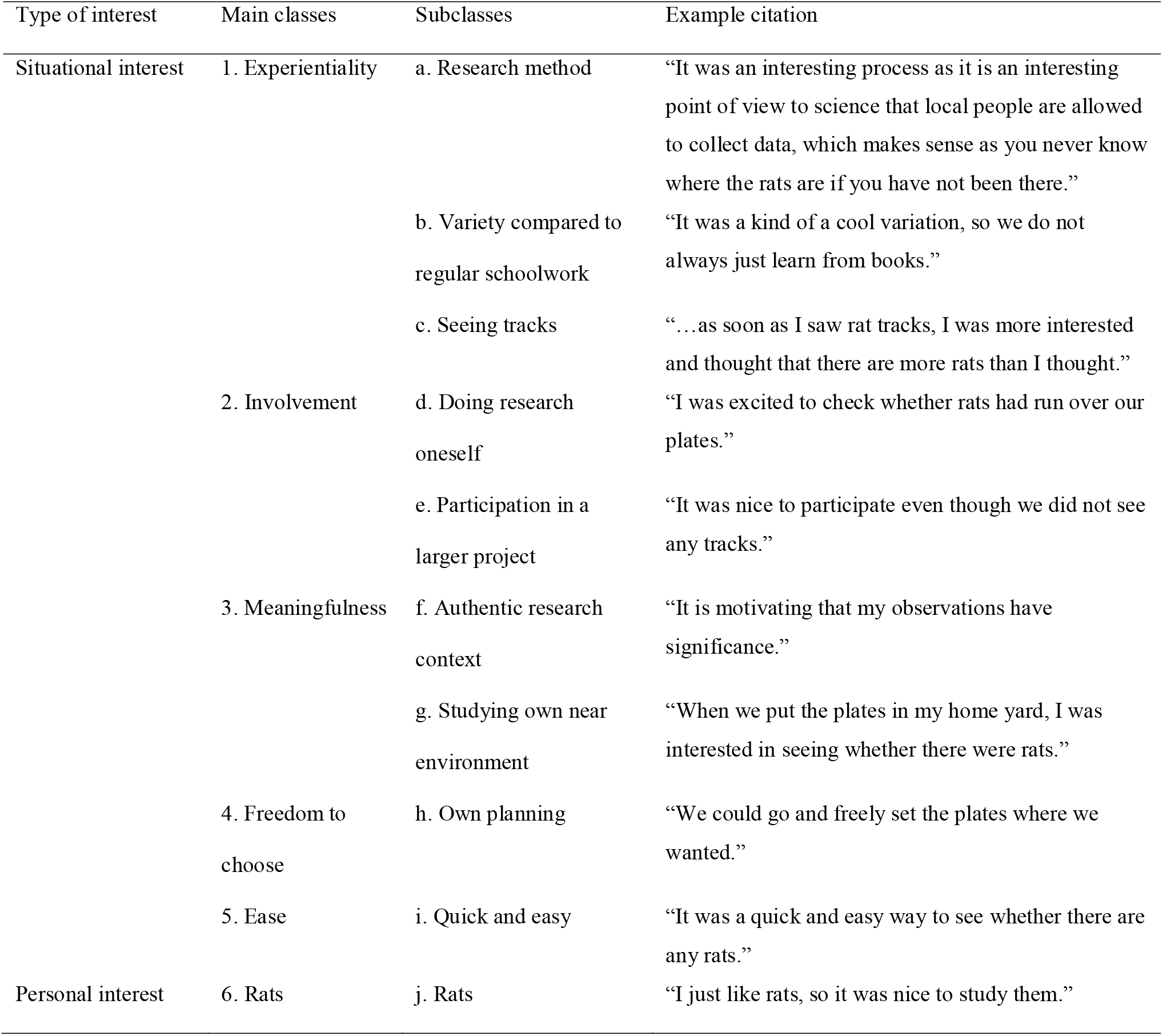
The characteristics of an urban rat citizen science project that increased interest in project participation

Situational interest was exhibited in five different main classes. *Experientiality* consisted of mentions where participation was something different compared to usual schoolwork or everyday life. *Involvement* consisted of comments either directed to specific parts of the research or in general participation to the larger research project. *Meaningfulness* was related to participation in an authentic research project or creating knowledge of one’s own near environment. Having *freedom to choo*se where to set the plates and the project being *easy to do* and straightforward were also seen as positive aspects. The only personal interest-related comments were made when students expressed their *positive attitudes toward rats* and how happy they were to study rats.

In comparison, the aspects that decreased interest related to the same issues. The most common aspect was a lack of reward: The project was seen as rather long and time-consuming compared to the results, and there were not enough results to analyze or failures in applying the research method itself. Lack of experientiality and meaningfulness were also mentioned: Students were disappointed when there were no rat tracks on the plates, or because they did not live within the study area, and could not study their own near environment.

#### Perceived learning during rat research

We classified students’ descriptions of their learning experiences with the framework of the revised Bloom’s taxonomy (Table 2). We did not observe all cognitive processes included in the taxonomy, such as analyzing or creating, because most of the mentions were on remembering and understanding. Remembering was mostly linked to factual knowledge, such as what rat tracks looked like, whether there were rats in their own near environment, and details on the natural history of rats:

> Student 6-2: “[I learned] How accommodating rats are and what strange environments they really can live in.”

**Table 2:**
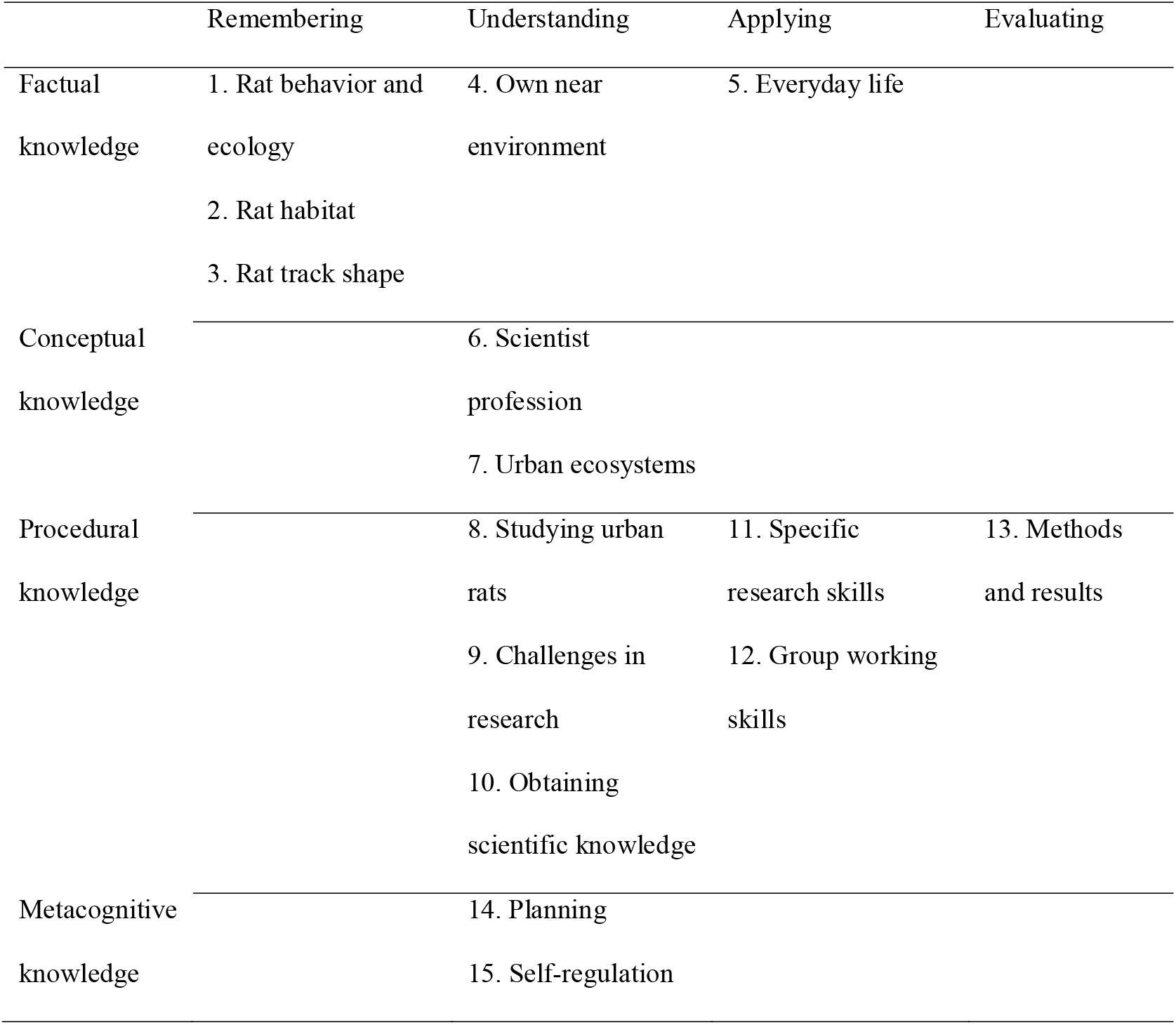
Self-described learning experiences by the students classified in the matrix of the revised Bloom’s taxonomy

Factual knowledge was also understood and applied specifically in the context of the students’ own near environment and everyday life.

> Student 4-3: “Before this project, I did not understand at all how many rats there actually are in Helsinki.† There was a lot of information that can be used in the future, like that I should not leave the trash can open.”

Described conceptual knowledge was limited to either understanding what urban ecosystems are and how they function or having a better grasp of what it is like to be a scientist. Most perceived learning was related to procedural knowledge on different cognitive process levels. While students described learning how to study rats in an urban environment, they also described different, broader topics, such as understanding how difficult it is to collect scientific data and how that knowledge actually is built.

> Student 9-2: “What we actually know does not just come from somewhere, but it needs to be properly researched.”

In addition, students described metacognitive processes such as understanding how they can plan the research and how they were able to work by following the instructions. More “higher”-level cognitive processes involved applying the research protocol in diverse authentic settings and being effective during group work. Furthermore, participation led the students to critically think about their accomplishments and the reliability of the data that they had collected.

> Student 9-1: “It would have been better to do this during summer, as now it snows, and you cannot leave the plates for four nights straight outside without them being covered in snow.”

In addition, one case of described learning did not fit the revised Bloom’s taxonomy because it was more related to attitudes toward performing research and an appreciation of the hard work needed to collect research data.

> Student 4-3: “… so many plates are needed that I kind of started to appreciate all the [research] data that are available on the Internet.”

### Feelings toward rats

In most of the interviews (six out of nine), students described negative feelings toward rats, while in the other three interviews, students described positive feelings toward rats (Table 3). When asked about the effect of participating in the project on their attitudes toward rats, seven students mentioned that they now have more positive attitudes.

> Student 5-1: “Rats are kind of stigmatized, but [that idea] kind of disappeared during this project as they do not seem to transmit diseases, so it does not really matter whether it is a squirrel or rat in my backyard.”

**Table 3:**
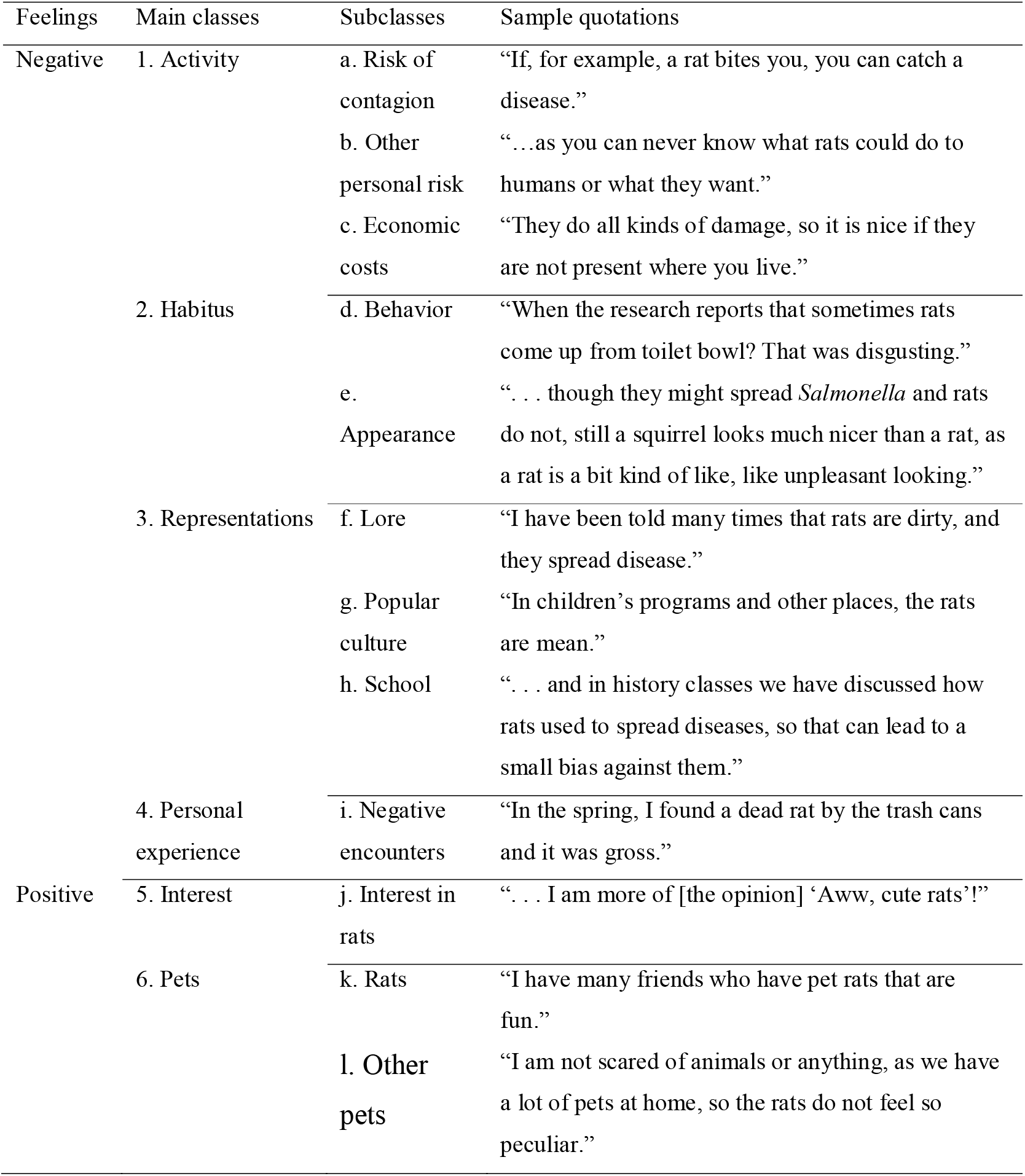
Classification of negative and positive feelings that the students described during the interviews

In contrast, one student mentioned having more negative feelings toward rats. They had not previously thought about rats and did not really understand that they are present everywhere in cities.

Rat activity was seen as problematic through the lens of perceived risks of contagion from rats or because of economic costs that the rats and their control pose on humans, including damage to infrastructure, such as chewed electric wires or broken trash cans. Other class of perceived risk was “another personal risk”, but this was not reflected in detail. The habitus of rats was seen as disgusting, whether it had to do with how rats move in the sewers or eat trash or because of their physical appearance. Rat representations in lore, popular culture, and school lessons were described as reasons why students had negative feelings toward rats. Students suggested that their aversion toward rats has been learned from society, or from school.

In contrast, the positive feelings described by two interviewees were linked to participation in the research; they had hoped to actually see rats and not only their tracks. Furthermore, one student suggested that having pets in general would lead to more positive feelings toward rats.

Though students commonly had negative feelings toward rats, none of the students suggested that these feelings would have prevented them from doing the study. The students mentioned that the research setting felt safe; thus, any worries they might have had were mitigated. Interestingly, when those students who had observed rat tracks in their own backyard or other near environment were asked whether that led to any actions, such as telling their parents, or whether they reflected on if that should affect their use of urban spaces, students did not describe any consequences as a result of observing rats.

## Discussion

We aimed to uncover how well participating in a citizen science project and surveying in the near environment the occurrence of an animal that usually elicits negative feelings can fit into a formal education setting. Our findings are encouraging from the point of view of the studied project and from a broader perspective of combining environmental education and citizen science. The students described multiple ways that participation led to an increase in their interest and resulted in diverse learning outcomes. Our findings are also in line with previous studies that highlight the potential of participating in citizen science projects during school time as a way of increasing personal interest in science (Hiller and Kitsantas 2014; Ruiz-Mallén et al. 2016).

### Authentic research context facilitates student interest

Students were curious about the urban rat research. As participation in the citizen science project provided a needed relief from usual schoolwork, it facilitated the awakening of situational interest, as suggested by Palmer (2009), in relation to inquiry-based learning. This experientiality remained throughout the research, as students were also interested in seeing the results of their study in a very practical way because they wanted to see the rat tracks on their plates. After the initial interest, a combination of two aspects was suggested to sustain interest. Firstly, authenticity of the project was linked to both involvement in a larger project and a broader meaning of the research work that the students conducted. Students were positively affected by the idea that what they were doing was not “in vain” but rather for a larger purpose as part of an actual research project. As found by Silva et al. (2002), this authenticity was not only a major motivational factor but also ensured that students would be more careful in working and generating reliable data. Secondly, contextuality was linked to the involvement and freedom in doing research and obtaining knowledge of their meaningful near environment. Our findings suggest that the students appreciated that they could use environments with which they were already familiar in their everyday lives as a study context. As such, citizen science projects in environments that are important for students can be very valuable contexts in facilitating learning (Pintrich, Marx, and Boyle 1993).

The factors that reduced interest in the research were most commonly the opposite of the factors that increased interest. While this is no surprise, one interesting tradeoff emerged in our analysis. While failure in the research or lack of rat tracks were seen as demotivating issues, they were still important learning experiences. Most of the students who suggested and had thought about improving the research setting or doing research in a different way—meaning they suggested learning higher cognitive skills—had failed in some way in their data collection. This suggests that errors in setting plates can lead a subset of students to a phenomenon similar to action adaptivity of error reaction as outlined by Dresel et al. (2013), which includes using (meta)cognitive activities to reflect on the error and thinking of improved working methods. It is noteworthy, though, that not all students who failed in data collection showed these deeper reflections. For most, the experience was described as demotivational.

### Learning with and about rats

Our findings revealed a diverse set of perceived learning that the students described in their interviews. While students mainly participated in the collection of data, as our project was contributory, this still led to potential substantial learning, although it has been suggested that learning outcomes are improved if participants are able to participate in many different aspects of a project (Zoellick, Nelson, and Schauffler 2012). As in previous research, students reported factual knowledge on study species and their near environment (Jordan et al. 2011; Mitchell et al. 2017; Silva et al. 2016) and procedural knowledge on used research protocols (Bonney, Cooper, et al. 2009; Bonney et al. 2016). In comparison, conceptual knowledge was not as well represented in students’ responses; for example, in a number of interviews students did not know what “urban ecosystem” meant. Contrary to results from previous studies (Bonney, Ballard, et al. 2009; Phillips et al. 2018), our participants suggested they had learned higher skills in planning and evaluating research settings, which might be because of participants’ broad autonomy in choosing sampling times and locations.

Rats evoke many kinds of feelings in students who study them. As in Kellert’s (1985) classification, rats were perceived as dangerous to humans, damaging pests, aesthetically suspect, and negatively linked through society and history to humans, though we did not find other Kellert’s classes. Both the negative cultural influences (Fox-Parrish and Jurin 2008; Prokop, Fančovičová, and Kubiatko 2009) and positive influences of having pets (Prokop and Tunnicliffe 2008) have been described in previous studies. The reasons for negative feelings toward rats were very similar to the reasons for negative feelings toward mice reported by Portuguese children (Almeida, Vasconcelos, and Strecht-Ribeiro 2014). We were not able to assess how attitudes changed during the project because we did not measure them prior to participation. Some students described themselves as having more positive attitudes toward rats, but it was more common that students had a simple acknowledgment of the presence of rats.

Interestingly, rats are ever-present in the urban areas, but students have rarely considered their presence. Rats occupy an interesting urban habitat, which does not correspond to what Jamie Lorimer (2015) describes as colonial imaginary of wilderness. The habitats of rats are very much spaces were humans are present, and, in fact, rats are there *because of* humans. Thus, rats do no occupy *wilderness*, but they are very much *wild* (Collard, Dempsey and Sundberg 2015). Nevertheless, there are encounters between students and rats, when rats run on the plates set by students, and students can observe these tracks, and thus opportunities for common worlds (Taylor and Pacini-Ketchabaw 2015, 2016).

While students were interested in knowing whether there were rats in their near environment, actual observations of rat presence did not lead to any substantial actions, such as reflections on their relationship to rats in an urban space or discussions with other people about rats. Thus, even though rats are an integral part of the urban ecosystem and demonstrably present in the nearest of near environments for students, the relationship between students and rats remains distant. Students described that they learned in school that rats spread diseases, which enforces the human-nature divide (Pacini-Ketchabaw and Nxumalo 2016); thus, a short project on actively creating knowledge about how humans and rats share urban space is not enough to counteract this. Thus, a deeper understanding of how humans and other animals share urban spaces and awareness of a near environment would require either more extensive research or more reflective research practices.

### Limitations of the study

The main limitation of our study was that we did not track actual learning outcomes but rather students’ perceptions and reports of what they had learned. As our study is based on the students’ self-described experiences, own interpretations, and self-reflection of what has happened, we are not able to assess the factual impact of our project in relation to their learning. Nevertheless, it is important to study the experiences of participants and what they perceive to learn from the project. It would be interesting to enrich this understanding of student experiences to observe actual research practices by shadowing students in the field while they are setting the plates.

The sample size was limited but the participants came from different age groups and different schools, so they were a good representation of the participants of the study overall. Even though the results might not be representative of all possible experiences, they describe well the experiences of the participants in the urban rat citizen science project. We do not know how much the motivation of the participants to take part in the interview biases our results. On the other hand, the most motivated students might be more willing to take part in interviews but it is also true that less motivated students might want to take part in interviews so they do not need to be in a biology class. Nevertheless, these are general problems related to the research interviews; therefore, any interview-based study is susceptible to these biases.

Our theoretical framework was not equipped to explore affective and emotional domains of learning about animals. Prior research suggests this is a very important part of learning with more-than-humans (Taylor, and Pacini-Ketchabaw, 2015; Boileau, and Russell 2018; Rautio et al. 2017; Lloro-Bidart 2018). Consequently, our project aims to delve deeper into the affective reactions to rats in further studies.

Notwithstanding the limitations in our study, we had similar results to many previous studies (Jordan et al. 2011; Mitchell et al. 2017; Silva et al. 2016; Bonney et al. 2016; Phillips et al. 2018). They suggest that citizen science projects can be a valuable tool in the palette of environmental education. Our results also suggest that students can learn higher skills in evaluation and planning research (Bonney, Ballard, et al. 2009; Phillips et al. 2018) and these student perceptions should be further studied by actually tracking learning outcomes.

### Implications for citizen science practitioners

There is commonly a potential conflict in citizen science projects because participants have different expectations from the project than the scientists running the project (Shirk et al. 2012; Zoellick, Nelson, and Schauffler 2012). This is understandable as scientists wish to collect as much reliable data as possible, whereas participants look for personal experiences (Rotman et al. 2012). Based on our results, we have evidence that the HURP is a citizen science project that provides learning opportunities for participating students. The clear and straightforward nature of our citizen science project was one of the advantages because students described the ease of participation as one of the characteristics that increased their interest in participating in the project.

The role of guiding and mentoring participation is emphasized in citizen science projects. As such, school groups are beneficial from the perspective of professional scientists because student participation is facilitated by their teachers. This provides helping hands for the project, as well-trained teachers are an asset in ensuring participation and providing support to participants.

Nevertheless, this creates an additional level of requirements for the citizen science projects because the projects need to be aligned to the school curricula. However, with a well-designed project, this can be a valuable asset because one of the perceived benefits of citizen science is learning, which is also the aim of school curricula.

### Implications for school practice

Our results support the idea that citizen science can be a valuable part of formal education in incorporating authentic research practices to everyday classroom practice. Furthermore, participating in a study coordinated by outside (i.e., academic) partners is seen as more valuable than experimental work done in school “for the school’s sake.” Especially in the cases where data collection occurs outside the school setting and even in the near environment for students, the motivational aspects of citizen science projects are significant.

Our study highlights some critical points that need to be considered for participation in citizen science projects to be successful. Firstly, the aims of teachers and students need to be aligned with the researchers who are running the project. In an optimal case, the citizen science project can fit perfectly to a curriculum and provide a ready-made package for teachers to use. Secondly, teachers cannot be passive participants in the project, but they need to react to the experiences and outcomes of student participation. We suggest that there is a need for discussion in the classroom on the objectives, problems related to, and eventually experiences during the citizen science project. While some students might acquire meaningful learning experiences just by participating, a chance to reflect on the project in a classroom would provide many more opportunities for learning. Thirdly, analysis of the data would provide further opportunities for learning; a collaborative or co-created citizen science project could be even more valuable from an educational point-of-view.

Nevertheless, there is a trade-off on how much course time teachers can dedicate to a citizen science project and how deeply students can become involved (Silva et al. 2016). We suggest that our approach can enable teachers to participate because it does not take much time, but this comes with a possible trade-off for more shallow learning experiences.

## Conclusions

Our case study of participation in a larger citizen science project suggests that studying one’s own near environment can be meaningful and experiential for students. Furthermore, autonomy and freedom in choosing study sites increased their described interest in the participation. Students described diverse perceived learning, including factual, conceptual, procedural, and metacognitive knowledge. In general, conceptual knowledge was less common than in previous studies of learning in citizen science projects, whereas procedural knowledge seems to have been more common.

For several reasons, students described negative attitudes toward rats. These included rat habitus and behavior, such as rats moving around in garbage and in sewers, but also cultural or personal experience, such as learning about rats in school lessons. Our results also suggest that studying a species that is commonly perceived as an unpleasant urban animal can have the potential to reduce negative attitudes towards the species.

## Acknowledgements

The authors wish to thank participating schools, teachers and students and Nina V. Nygren and Pauliina Rautio for discussions on human-animal relations and Mary Lukkonen and Marlene Boemer on proof-reading the manuscript. T.A. has been funded by the Maj and Tor Nessling Foundation and the City of Helsinki research grants. The authors report no potential conflict of interests.

## Biographical note

*Tuomas Aivelo* is a postdoctoral researcher in Organismal and Evolutionary Biology, University of Helsinki and coordinator of the Helsinki Urban Rat Project. Aivelo studies both ecology and evolution of animals, with an emphasis on parasites and is involved in biology education. He is also keen on science outreach and writes a popular blog on parasites in collaboration with biology and geography teachers in Finland.

*Suvi Huovelin* is a recently graduated biology and geography teacher in Helsinki, Finland. This article outlines the main findings of her Master of Science thesis.

